# Hydrothermal vent fauna of the Galápagos Rift: Updated species list with new records

**DOI:** 10.1101/2023.11.28.568903

**Authors:** Chong Chen, John W. Jamieson, Verena Tunnicliffe

## Abstract

The sighting of giant bivalves and tubeworms at the Rose Garden vent field on the Galápagos Rift in 1977 marked the discovery of hydrothermal vents, a turning point for modern biology. The following decade saw a flurry of taxonomic descriptions of vent endemic species from the first vents. With the finding of high-temperature ‘black smokers’ on the East Pacific Rise, exploration shifted away from Galápagos. A faunal list of Galápagos vents with 65 species was published in 1991, then updated to 74 species in 2006. Since then, few expeditions returned to the Galápagos Rift. Here, we revisited several Galápagos vents including recently confirmed high-temperature sites and inactive sulfide mounds. From our collecting efforts and observations, as well as revisions from the literature, we update the faunal list to 92 species including 15 new records, restricted to obvious vent associates. Accurate regional faunal lists are important for understanding the biogeography of vent fauna, and our list will also be valuable for setting management strategies.

## Introduction

The discovery of hydrothermal vents themselves on the eastern Galápagos Rift in 1977 was not really a surprise, as geologists had predicted their presence from the missing heat measured near ridge axes (Sclater and Klitgord 1973) and warm buoyant plumes collected by a towed vehicle (Weiss et al. 1977). But nobody was prepared for the first contact with its bizarre inhabitants at a supposedly nutrient-deficient deep seabed two and a half kilometres below the surface – dense aggregations of giant clams and mussels, worms living in metre-long white tubes with red plumes swaying in shimmering water, and all other animals living with them (Corliss et al. 1979). Starting with the description of the giant vent clam *Turneroconcha magnifica* and the bythograeid vent crab *Bythograea thermydron* (Boss and Turner 1980; Williams 1980) followed by the giant tubeworm *Riftia pachyptila* and the discovery of chemosymbiosis (Cavanaugh et al. 1981; Jones 1981), biologists began to tease apart the taxonomic affinities and evolutionary origins of these creatures (Hessler and Smithey 1983).

Within a decade of its discovery, nearly all animals found in the original diffuse flow Galápagos Rift vents, now known as the Rose Garden vent field, were described. An early faunal list in 1991 included 65 species (Tunnicliffe 1991; Tunnicliffe 1992), and another in 2006 listed 74 (Desbruyères et al. 2006). With the discovery of ‘black smoker’ chimneys spewing out high-temperature fluids in other systems such as East Pacific Rise (EPR) and Juan de Fuca Ridge (Tunnicliffe et al. 1985; Desbruyères and Laubier 1986), exploration shifted away from Galápagos Rift where vigorous venting was apparently lacking. An expedition in 2002 found Rose Garden had been buried under fresh basaltic lava flows, and the communities had been largely wiped out except some recent settlers on nearby low-temperature venting from cracks in an area named Rosebud (Shank et al. 2003). The 2002 expedition also found signals for more vents east of Rose Garden on the eastern rift, plus a vent at the western Galápagos Rift near the Galápagos Islands. Between 2005-2006, towed-camera surveys in the western Galápagos Rift confirmed the first high-temperature chimneys (Haymon et al. 2008). Though these sites provide likely grounds for new records and subsequent research cruises have visited some of these areas using underwater vehicles (Shank et al. 2012; Raineault et al. 2016), no faunal updates have been published to date.

From October to November 2023, we were able to revisit the Galápagos Rift vents on-board the Schmidt Ocean Institute’s R/V *Falkor (too)* during the research cruise FKt231024. One aim of the cruise was to investigate the distributions of animal communities associated with both active and inactive vents. Here, we revise the faunal list of Galápagos Rift hydrothermal vents based on the literature since the last compilation (Desbruyères et al. 2006) plus new findings from our research expedition, in order to present all reliable distribution records from vents in this region.

## Materials and Methods

During R/V *Falkor (too)* cruise Fkt231024, we visited several hydrothermal vent fields on the Galápagos Rift using the remotely operated vehicle (ROV) *SuBastian*. These included Rose Garden / Rosebud (0.81°N, 86.22°W, 2450-2550 m deep; dive #603) and Tempus Fugit (0.77°N, 85.91-93°W, 2500-2560 m; dive #606-607, 609) on the eastern Galápagos Rift (Shank et al. 2012; Raineault et al. 2016); as well as Iguanas-Pinguinos (2.10°N, 91.89-94°W, 1650-1700 m; dive #611-613) and Tortugas, a newly-discovered active vent field on the western edge of the East Los Huellos Caldera (0.95°N, 90.53-56°W, 1500-1600 m; dive #614) on the western Galápagos Rift (Haymon et al. 2008). An overview map of the study area is presented in Figure 1.

**Fig. 1.**
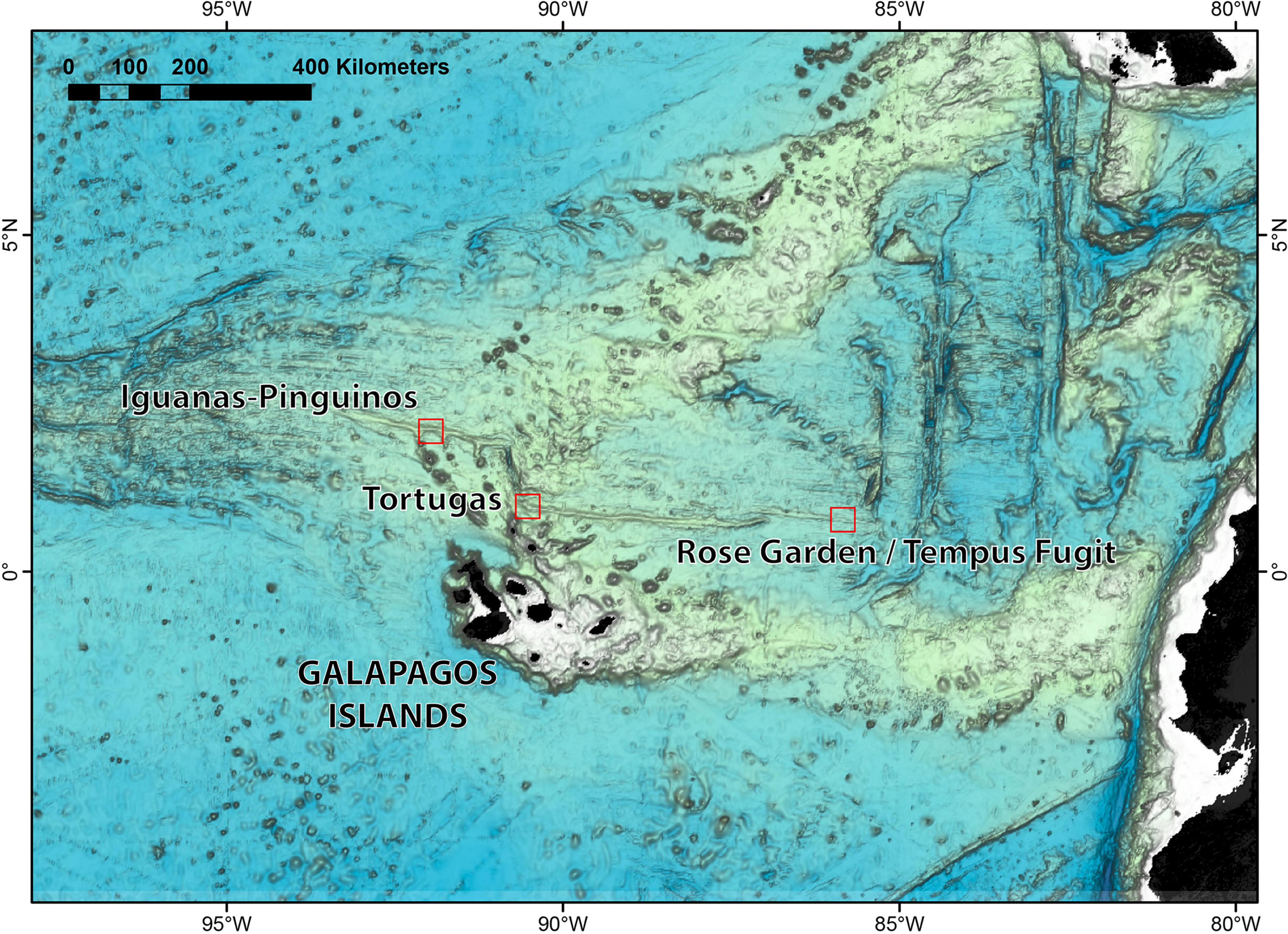
Map of the study area, showing the location of each hydrothermal vent field visited during R/V *Falkor (too)* research cruise Fkt231024.

We used video and screengrabs from a 4K ultra-HD video camera (SULIS Subsea Z70; resolution 3840 x 2160 pixels) on the ROV *SuBastian* for seafloor imaging, which allowed up to 12x zoom for close-up observations of even smaller animals. Animals were collected using either a seven-function manipulator arm (Schilling Robotics TITAN 4) or a suction sampler mounted on ROV *SuBastian*. Upon recovery on-board, animals were sorted in cold (4°C) seawater, cleaned with a brush, and photographed using a Canon EOS 5Ds R digital single-lens reflex camera equipped with a Canon EF 100 mm F2.8L MACRO IS USM macro lens. Most new records are based on collected specimens, but some larger fauna were identified using close-up imagery. For peltospirid gastropods, which exhibited considerable morphological variability compared to specimens known from the EPR (McLean 1989a), the barcoding fragment of the mitochondrial cytochrome *c* oxidase subunit I (COI) gene was amplified and sequenced using the universal primer pairs HCO2198-LCO1490 (Folmer et al. 1994) following a published protocol (Chen et al. 2018) to confirm their identities. The new sequences were deposited on GenBank (PP000825-PP000827) and were compared with existing EPR sequences using the search function and the built-in pairwise distance calculator of NCBI BLAST.

Previously published faunal lists for the Galápagos Rift vents (Tunnicliffe 1991; Tunnicliffe 1992; Desbruyères et al. 2006) were examined for taxonomic status using both the World Register of Marine Species (WoRMS Editorial Board 2023) and primary literature. We aimed to remove erroneous records and to ensure the list only includes those species that rely strongly on the vent environment. To eliminate erroneous records, occurrence records at the Galápagos Rift were checked against the original descriptions and subsequent works on each species; geographic distribution of those species found in other hydrothermal systems were also recorded (see Table 1). New species described since the publication of the previous lists were checked in Google Scholar using search terms “Galapagos AND hydrothermal AND new species”. New records from our present study are added to this ‘historical’ list.

**Table 1.**
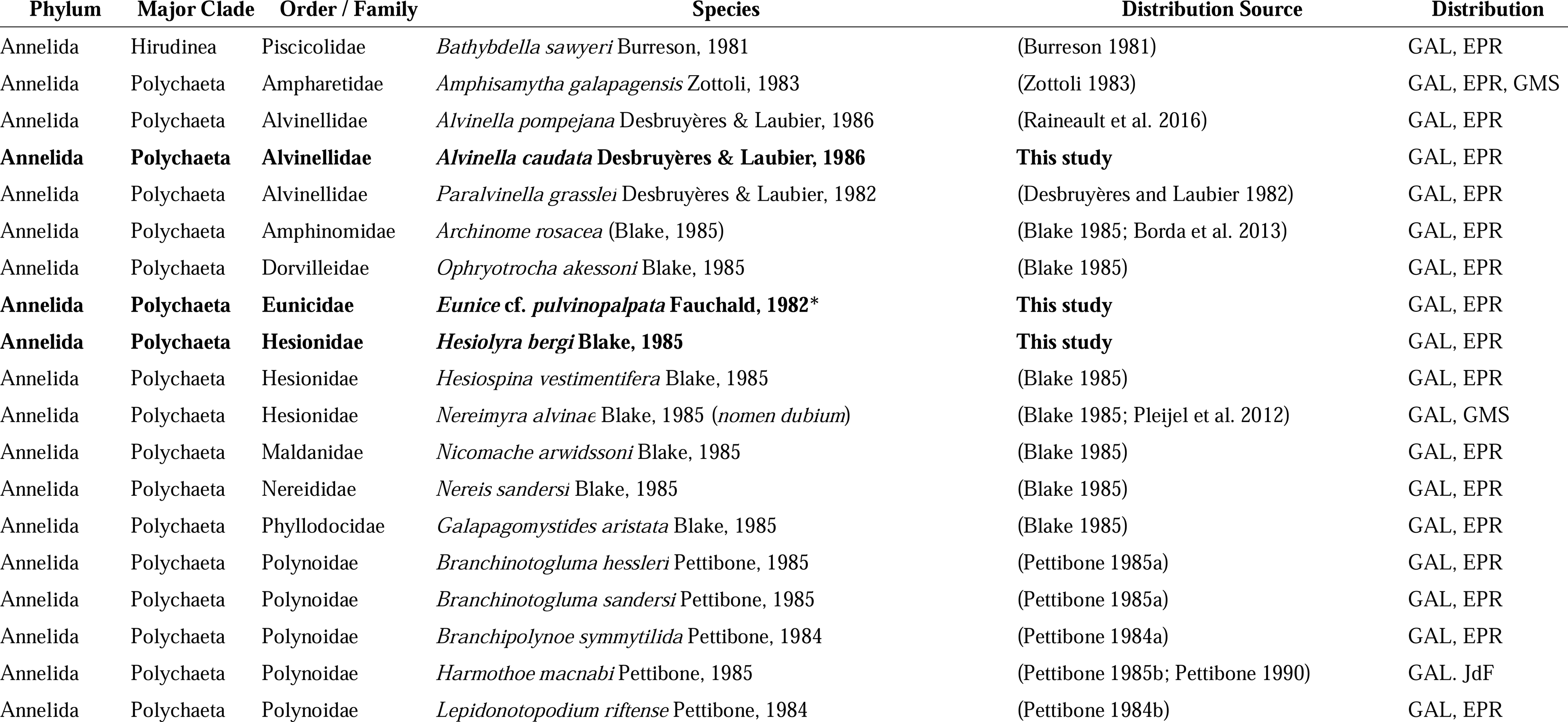
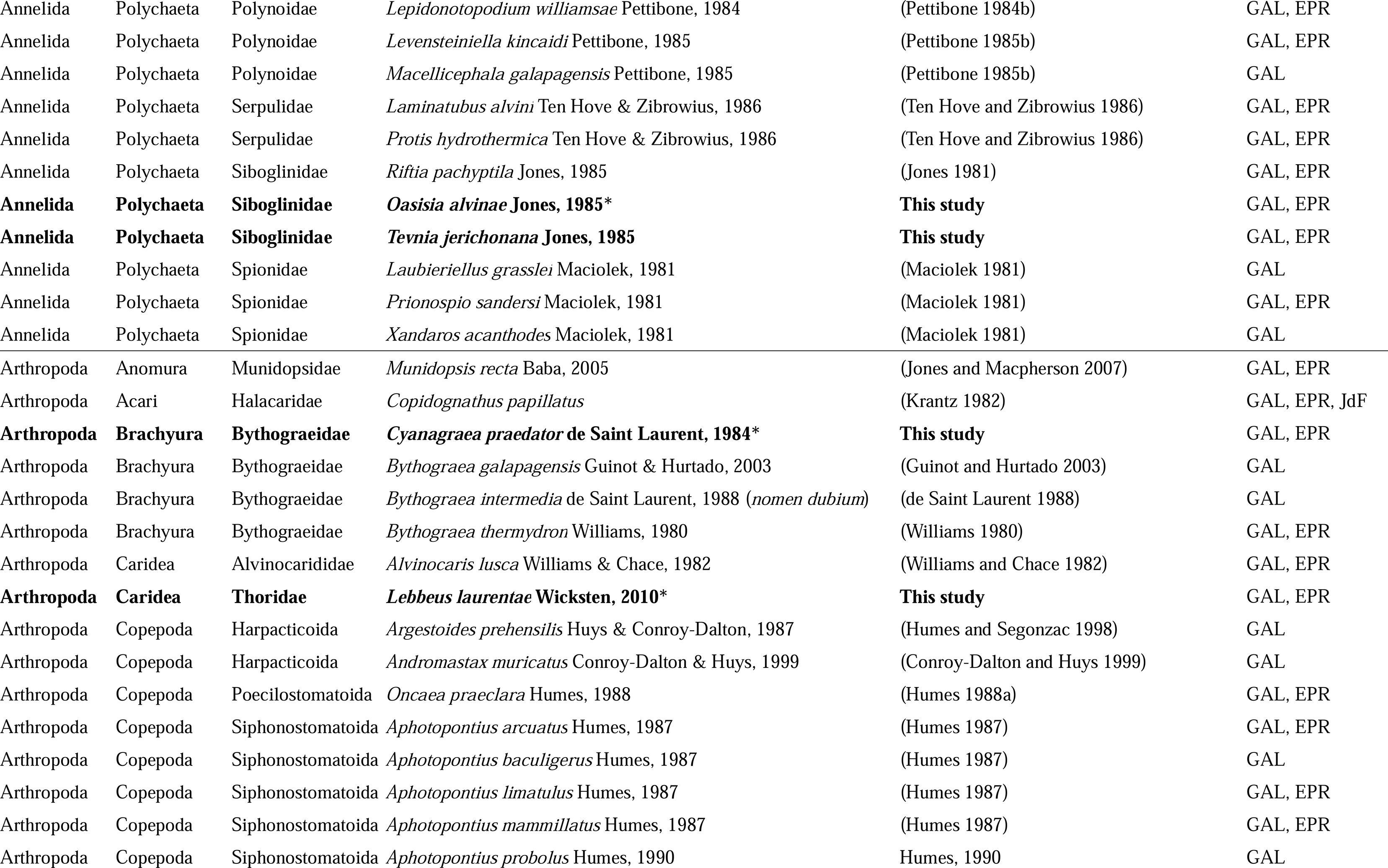
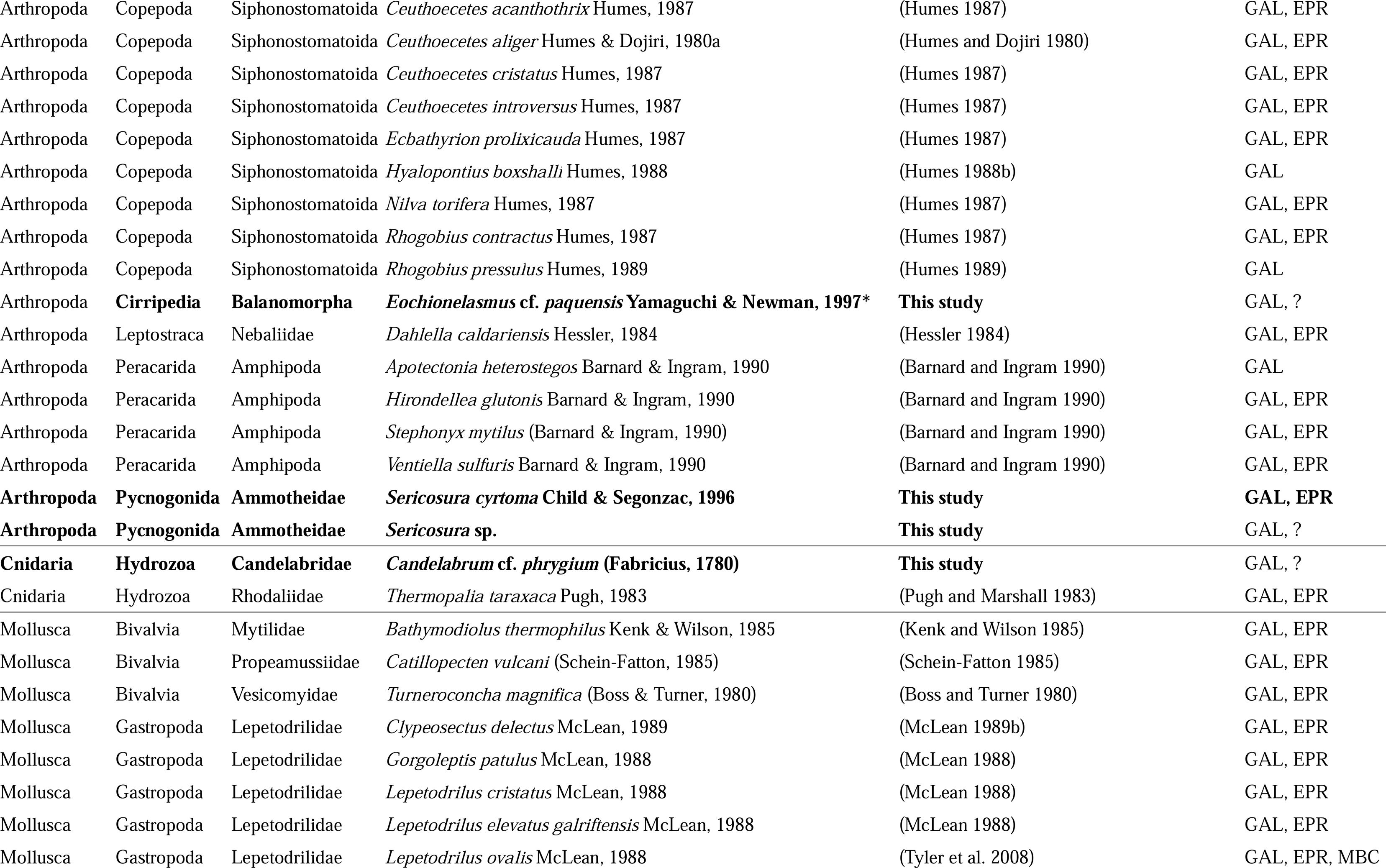
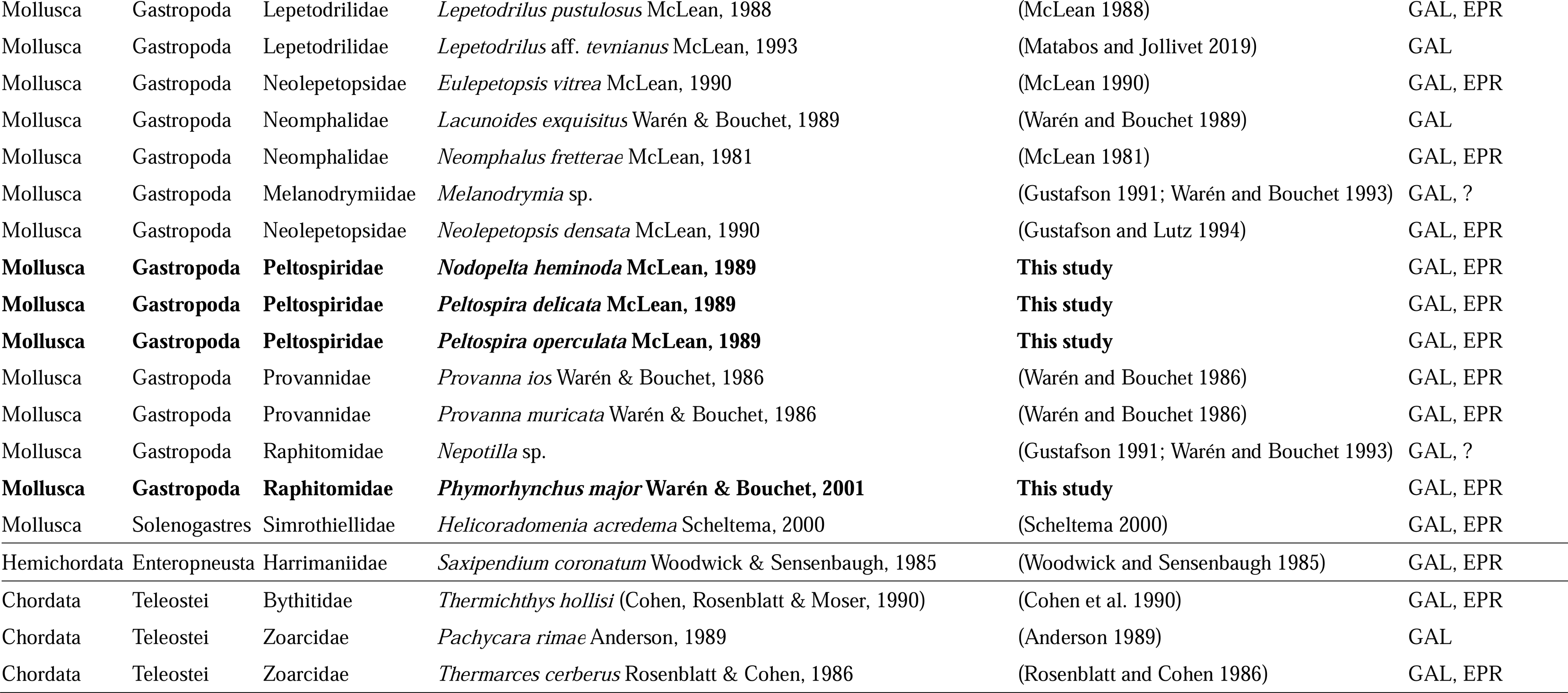
List of species recorded at Galápagos Rift hydrothermal vent systems. Species newly recorded in this study are listed in bold, species records from imagery only are denoted with asterisks and their species-level identification should be considered tentative. Abbreviations: GAL = Galápagos Rift; EPR = East Pacific Rise; GMS = Guaymas Basin; MBC = Monterey Bay, California; JdF = Juan de Fuca Ridge. “?” in distribution means occurrence outside Galápagos Rift remains uncertain.

## Results and Discussion

### Overview of vent fields visited

In the eastern Galápagos Rift, the Rose Garden / Rosebud area was covered by fresh basaltic lava flow and devoid of living vent fauna. This confirms the finding from a 2011 cruise that another eruption event between 2005 and 2011 had eliminated fauna in this field (Shank et al. 2012). Furthermore, we revisited a serpulid worm colony found in 2015 (Raineault et al. 2016) in case living vent fauna persisted (0.8049°N, 86.2194°W, 2447 m deep), but only found decaying serpulid tubes and dissolving mussel shell debris. As such, venting at Rose Garden has likely ceased – although we did not visit the location of the East of Eden field. In the nearby Tempus Fugit field (Raineault et al. 2016), we found that venting at the previously known main diffuse flow site (0.7700°N, 85.9114°W, 2561 m deep) had waned, with few living vesicomyid clams and *Riftia* tubeworms. Nevertheless, we found a new diffuse flow vent nearby (0.7712°N, 85.9236°W, 2602 m deep; ‘Walking Dead’ vent). We also revisited the active chimney (Raineault et al. 2016) at the western end of Tempus Fugit (0.7712°N, 85.9332°W, 2514 m deep; ‘Zombie’ vent) and confirmed high-temperature (>200°C; measured with the ROV temperature probe) venting there. A number of dead spires or inactive mounds were found around the Zombie vent and were also surveyed.

Shifting to the western Galápagos Rift, we revisited all three vent sites in the Iguanas-Pinguinos vent field (Haymon et al. 2008; Raineault et al. 2016), including Iguanas West (2.0992°N, 91.9053°W, 1670 m), Iguanas East (2.1050°N, 91.9378°W, 1670 m), and Pinguinos (2.0993°N, 91.9052°W, 1670 m). We confirmed chimney structures associated with vigorous venting of high-temperature fluid at all three locations. At East Los Huellos Caldera, where only plume signals were known (Haymon et al. 2008), we discovered active venting associated with chemosynthetic communities. This included both diffuse venting areas dominated by mussels and vesicomyid clams (0.9546°N, 90.5566°W, 1590 m) and active chimney complexes with high-temperature (>250°C) venting (0.9543°N, 90.5613°W, 1520 m).

### Revising the existing faunal list

The most recent faunal list of the Galápagos Rift vents (Desbruyères et al. 2006) included a total of 74 species. Of these, two orbiniid annelid species including *Orbiniella aciculata* and *Scoloplos ehlersi* were erroneously included in the list, as the author clearly states these were collected from box cores deployed near the Galápagos Rift but were not from the vent community (Blake 1985). Here, we further remove the lysianassoid amphipod *Abyssorchomene abyssorum* on the grounds that it is a globally distributed deep-sea species found in non-chemosynthetic seafloor and that its vent record is based on a single specimen that may have been a by-catch (Barnard and Ingram 1990). Similarly, we took out the abyssal grenadier *Coryphaenoides armatus* since it is merely an occasional visitor to vents from the surrounding deep sea. Though there are two species of dubious taxonomic status – the hesionid polychaete *Nereimyra alvinae* with poorly preserved types (Pleijel et al. 2012) and the crab *Bythograea intermedia* described from megalopa and juveniles only (de Saint Laurent 1988) – we have kept them, pending future taxonomic revision. The melanodrymid snail *Melanodrymia* sp. and the raphitomid snail *Nepotilla* sp. were initially reported in a conference abstract (Gustafson 1991) and then included in a gastropod faunal list by taxonomic experts (Warén and Bouchet 1993). Though their species-level identification remains unclear, they remain on the list pending more taxonomic information. Moalic et al. (2012) supplemented the list in Desbruyères et al. (2006) to report 83 taxa. In addition to the annelids above, occasional visitors and a double record of *Thermichthys hollisi*, we removed species we could not verify such as polychaetes only known from Juan de Fuca/Gorda Ridges, *Bythograea microps,* and *Aphotopontius acanthinus.* There remained 75 species.

Since the 2006 list was published, three additional species have been recorded from Galápagos Rift in the published literature. The first is the squat lobster *Munidopsis recta* Baba, 2005 that was confirmed as a Galápagos record by Jones and Macpherson (2007) using COI sequencing. The second species is the Pompeii worm *Alvinella pompejana*, visually confirmed from Tempus Fugit vent field in 2010, but not sampled (Raineault et al. 2016). The third species is *Lepetodrilus* aff. *tevnianus* Galápagos *sensu* Matabos and Jollivet (2019), morphologically resembling *Lepetodrilus tevnianus* found on the EPR vents but is a genetically distinct lineage considered to represent an undescribed species (Matabos and Jollivet 2019). Altogether, these bring the historical species occurrence record to 77 species.

### New records

From our observations and collections during the 2023 cruise, we encountered a total of 15 species that are clearly associated with the chemosynthetic ecosystem and not previously recorded from Galápagos Rift vents (Tunnicliffe 1992; Desbruyères et al. 2006). Table 1 lists our updated full faunal list comprising 92 species, with our new records shown in bold. Figure 2 presents key *in situ* screengrabs including records based on species clearly identifiable from imagery, while figure 3 shows photographs of specimens collected. In the following paragraphs, we provide more details on our newly recorded species.

**Fig. 2.**
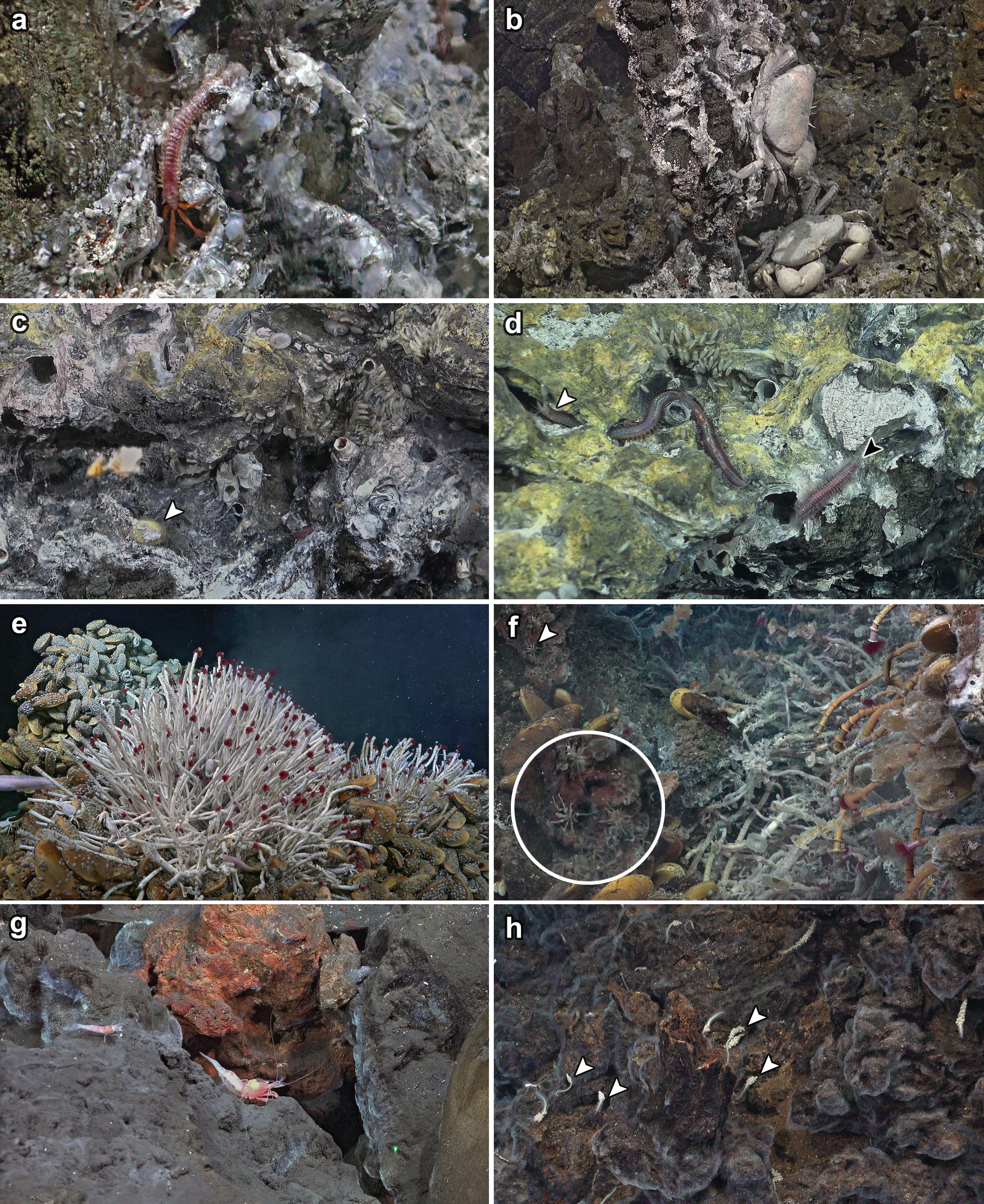
*In situ* imagery of Galápagos Rift vents captured by screengrabs of the 4K video camera in the present study: **a** the alvinellid worm *Alvinella caudata* on chimney wall of Zombie vent, Tempus Fugit; **b** two individuals of the bythograeid crab *Cyanagraea praedator*, Zombie vent, Tempus Fugit; **c** a living individual of *Nodopelta heminoda* (white arrow), Zombie vent, Tempus Fugit; **d** the polychaete worms *Eunice* cf. *pulvinopalpata* (white arrow) and *Hesiolyra bergi* (black arrow), Zombie vent, Tempus Fugit; **e** a bouquet of *Tevnia jerichonana* tubeworms at a peripheral diffuse flow at West Iguanas, Iguanas-Pinguinos; **f** a cluster of *Oasisia alvinae* tubeworms at a diffuse flow site in East Los Huellos Caldera and several individuals of *Sericosura* pycnogonids nearby (white arrow and enlarged in inset; likely a mix of both species on Table 1); **g** *Lebbeus laurentae* (larger shrimp on the right) seen with *Alvinocaris lusca* at the base of the active chimney complex at West Iguanas, Iguanas-Pinguinos; **h** several hydrozoan *Candelabrum* cf. *phrygium* near a low-temperature vent at West Iguanas, Iguanas-Pinguinos (representative individuals indicated by white arrows).

**Fig 3.**
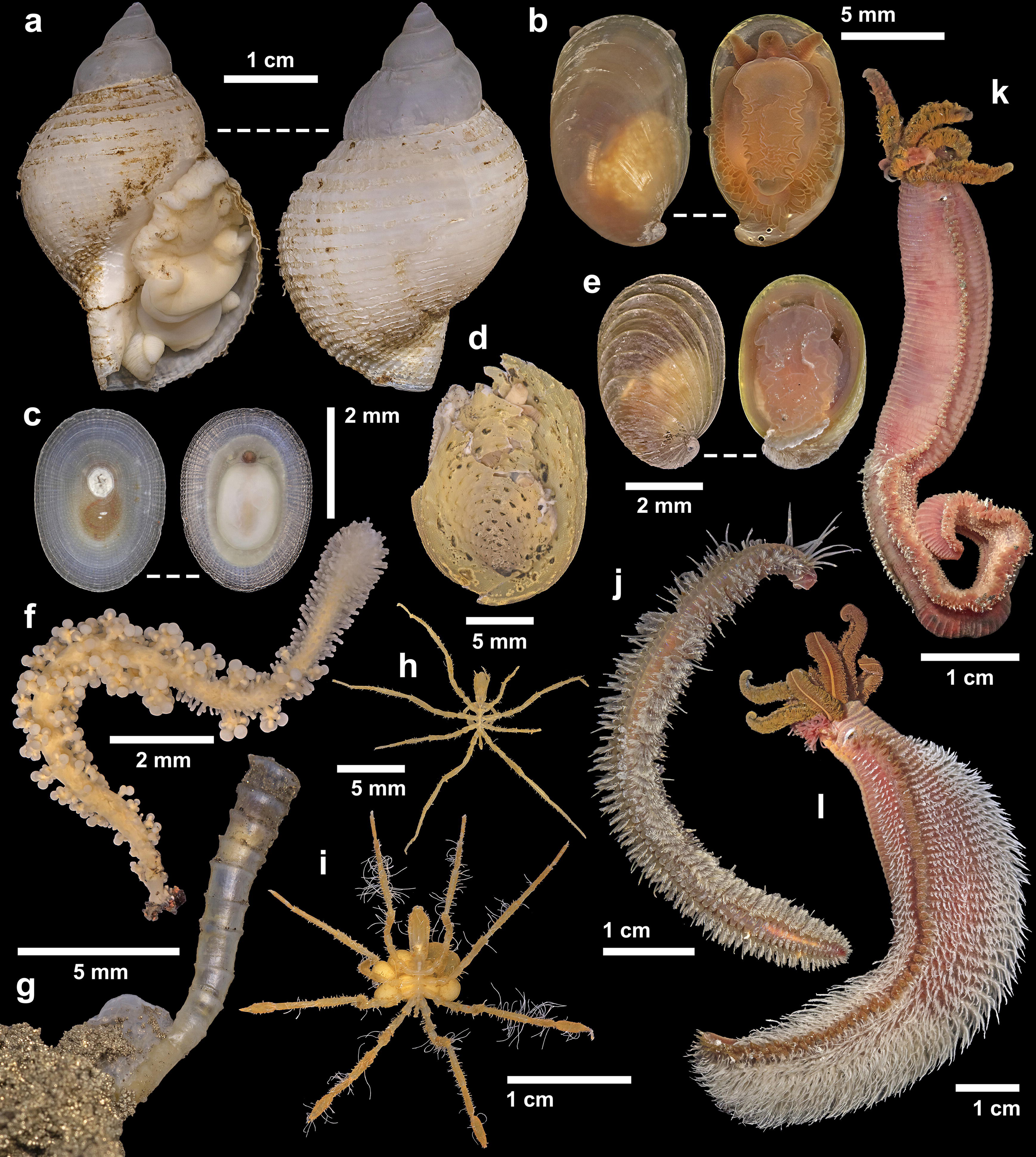
Specimens collected from Galápagos Rift vents in the present study. **a** *Phymorhynchus major*, Walking Dead diffuse flow vent, Tempus Fugit; **b** *Peltospira* sp., Zombie vent, Tempus Fugit; **c** *Neolepetopsis densata*, inactive spires near Zombie vent, Tempus Fugit; **d** *Nodopelta rigneae*, Zombie vent, Tempus Fugit; **e** young individual of *Peltospira delicata*, Zombie vent, Tempus Fugit; **f** *Candelabrum* cf. *phrygium*, West Iguanas, Iguanas-Pinguinos; **g** a juvenile individual of *Tevnia jerichonana* from Zombie vent, Tempus Fugit; **h** *Sericosura* sp., diffuse flow at East Los Huellos Caldera; **i** *Sericosura cyrtoma*, diffuse flow at East Los Huellos Caldera; **j** *Hesiolyra bergi*, Zombie vent, Tempus Fugit; **k** *Alvinella caudata*, Zombie vent, Tempus Fugit; **l** *Alvinella pompejana*, Zombie vent, Tempus Fugit.

The alvinellid worm *Alvinella caudata* (Fig. 2a, 3k) was seen on active chimney walls at Tempus Fugit, Iguanas-Penguinos, and Tortugas. It co-occurred with *A. pompejana,* and both were collected together at Tempus Fugit; we show a specimen photo of *A. pompejana* (Fig. 3l) since this is the first time a specimen was collected from Galápagos Rift and serves as a confirmation of the previous record (Raineault et al. 2016). Also found on the same habitat was the hesionid worm *Hesiolyra bergi* (Fig. 2d, 3j), which occurred in aggregations on the chimneys; individuals were sometimes seen going into tubes of *Alvinella* worms. We give a tentative identification of *Eunice* cf. *pulvinopalpata*(Fauchald 1982) to a eunicid worm seen in the same habitat, but not collected (Fig. 2d). The bythograeid crab *Cyanagraea praedator* (Fig. 2b) was common on the active chimneys too, readily identified from images by the well-developed eye-stalk sockets and their large size (de Saint Laurent 1984). The association between *Cyanagraea* and *Alvinella* is also known from EPR vents, where the former is a predator of the latter (Desbruyères et al. 2006).

We also collected three species of peltospirid gastropods from the high-temperature Zombie vent at Tempus Fugit, including *Nodopelta heminoda* (Fig. 3d), *Peltospira delicata* (Fig. 3b), and *Peltospira operculata* (Fig. 3e). Although only one damaged specimen of *N. heminoda* could be collected, several individuals were seen near *Alvinella* tubes (Fig. 2c); the COI sequence of the collected specimen (GenBank PP000825) matched an existing mitogenome of the same species (GenBank BioProject PRJNA927338) with a pairwise identity of 99.82%. *Peltospira operculata* is also recorded based on a single young specimen (Fig. 3e) still displaying strongly ribbed shell sculpture that fades in adults (McLean 1989a). The spacing of its ribbing is wider than typical specimens from the EPR (McLean 1989a) but its COI sequence (GenBank PP000826) was closely comparable to five existing sequences (GenBank GU984275-GU984279) with pairwise identities between 99.19-99.68%, indicating this spacing is intraspecific variation. *Peltospira delicata*, recorded based on two collected adult specimens, was unusual in lacking clear spiral ridges on the body whorl (McLean 1989a). The COI sequence of the ethanol-preserved specimen (GenBank PP000827) matched an existing sequence of *P. delicata* (GenBank AY923931) with a pairwise identity of 99.85%. Though having a smooth adult shell is reminiscent of *Peltospira operculata*, all other external morphological features of our specimens such as the overall weaker coiling and the lack of operculum agree with identification as *P. delicata* (McLean 1989a; Warén and Bouchet 2001). This is indicative of a wider range of phenotypic variability in this species than previously known. Peltospirid snails were not seen on active chimneys in the western Galápagos Rift, but as we did not sample those sites, we may have missed them on video due to their small size.

At both diffuse flow areas and active chimney walls in Iguanas-Pinguinos and Tortugas we saw bouquets of the tubeworm *Tevnia jerichonana* (Fig. 2e), and a single juvenile specimen (Fig. 3g) was collected from the Zombie vent at Tempus Fugit where no adults could be seen. Only at the diffuse flow site at Tortugas, did we see a cluster of *Oasisia alvinae*. The only tubeworm known from previous explorations in the eastern Galápagos Rift was *Riftia pachyptila* (Corliss et al. 1979; Jones 1981; Raineault et al. 2016), which also occurred in both diffuse flow sites in Tempus Fugit (but in lower abundance than previous expeditions due to waning activity there). Conversely, at the western Galápagos Rift we did not see any sign of *Riftia*. At Tortugas, we found two species of the pycnogonid genus *Sericosura* in abundance around diffuse flows (Fig. 2f). One species with seven-segmented palps was readily identifiable as *Sericosura cyrtoma* (Fig. 3h), but the other (Fig. 3i) with nine-segmented palps did not match any described eastern Pacific congeners (Child 1987; Child and Segonzac 1996; Wang et al. 2013) and may represent an undescribed species. Though we did not find pycnogonids in the eastern Galápagos Rift, a previous cruise reported seeing pycnogonids there (Raineault et al. 2016), likely also *Sericosura*.

The raphitomid snail *Phymorhynchus* was often seen in the periphery zone of all vent fields we visited. Initially, *Phymorhynchus* from the Galápagos Rift was considered to be conspecific with those on the EPR (Warén and Bouchet 1989), but this distribution record was not mentioned when *P. major* was formally named based on only EPR material (Warén and Bouchet 2001). Here, we collected a specimen (Fig. 3a) and confirm the presence of *P. major* in the Galápagos. Though not seen on our expedition, we note that a recent expedition also on R/V *Falkor (too)* (Fkt230812) encountered dense coverage of a vent barnacle tentatively identified as *Eochionelasmus* cf. *paquensis* (Hiromi K. Watanabe, pers. comm.) at a vent site named Sendero del Cangrejo (2.53°N, 94.33°W, 2490 m deep). This species is added to our list based on imagery shown on an openly available YouTube stream of ROV *SuBastian* dive #573 at this site (Schmidt Ocean Institute 2023).

We also saw several individuals of *Lebbeus* co-occurring with *Alvinocaris lusca* on the vent periphery only in the West Iguanas vent (Fig. 2g). Though the *Lebbeus* was not collected our imagery provided sufficient resolution for its tentative identification as *L. laurentae* based on external morphology (Komai et al. 2012). Numerous individuals of the hydrozoan *Candelabrum* were seen also near the periphery of West Iguanas (Fig. 2h). The collected individual (Fig. 3f) was morphologically similar to *Candelabrum phrygium* which has a pan-arctic distribution and also known from Mid-Atlantic Ridge vents (Segonzac and Vervoort 1995). As Galápagos is far from its known range, we consider it likely to be a distinct species and tentatively identified it as *C.* cf. *phrygium*. Further away from high-temperature venting, we found many individuals of the true limpet *Neolepetopsis densata* on inactive chimneys near Zombie vent near Tempus Fugit (Fig. 3c). Although Gustafson and Lutz (1994) published a record for *N. densata* from an inactive mound on the Galápagos Rift, the validity of this was questionable as the figure captions listed the illustrated specimens as from Galápagos but the same figures were cited in the main text as specimens from 9-10°N on the EPR. Our present finding serves to confirm their Galápagos record. To our knowledge this is the only Galápagos Rift species likely restricted to inactive chimneys, a distribution pattern typical for genus *Neolepetopsis* (McLean 1990; Chen et al. 2021).

During our exploration we also saw a number of animals typical of non-chemosynthetic seafloor environments within proximity to vents, such as the Pacific white skate *Bathyraja spinosissima* known to incubate egg cases at Galápagos Rift vents (Salinas-de-León et al. 2018), the octopus *Graneledone* (likely an undescribed species, Janet Voight pers. comm.) (Desbruyères et al. 2006), and some encrusting demosponges. We did not include them in our list due to the likely incidental nature of their presence in or near the chemosynthetic ecosystem.

We note that a limitation of our study is that some new records such as the tubeworm *Oasisia alvinae* or the vent crab *Cyanagraea praedator* were not collected and identified based on imagery data only, precluding future genetic studies. Though these species are easily identified from external morphology based on our current understanding with just one species in their respective genera in the eastern Pacific vents, we cannot rule out the presence of cryptic species specific to the Galápagos Rift. Previous studies in annelids and gastropods have highlighted the presence of genetic barriers and population subdivisions between the Galápagos Rift and the EPR (Hurtado et al. 2004; Matabos and Jollivet 2019). This includes cryptic species that are separated across the two ridge systems; for example the limpet *Lepetodrilus elevatus* is known to consist of at least four cryptic genetic lineages across the eastern Pacific vents that are tentatively treated as one species (Matabos and Jollivet 2019).

Forty-five years after the discoveries at Rose Garden, there still are vent communities within 35 km of the original site. As the most easterly extension of the east/southeast Pacific biogeographic region, these vents may be both a population sink and a source of novel genetic diversity. Given the location in international waters near the large Galápagos Mounds sulphide deposits, consideration of protections such as Ecologically and Biologically Sensitive Area (EBSA) designation is warranted. The abundant active and inactive chimneys of the Iguanas-Pinguinos and Tortugas sites are testament to long-term hydrothermalism that has supported vent communities and diversification of the fauna. Currently, at least 14 species (not including *nomen dubium*) are known only from the Galápagos Rift (Table 1), an endemism proportion of 15%; another five species whose endemism is uncertain. While not high compared to the endemism among western Pacific vent systems (Tunnicliffe et al. 2023), as there are no geographic barriers separating the Rift from EPR, specific environmental conditions may foster the endemics. For example, sustained venting over numerous large chimneys may foster population maintenance compared to the high turnover at EPR vents (Gollner et al. 2017). Further collecting and molecular work are needed to investigate the biogeographic relationships between Galápagos Rift and the EPR in finer detail. Accurate species lists and occurrence data can reveal key processes driving the biogeographic patterns and evolution of hydrothermal vent fauna in general (Giguère and Tunnicliffe 2021; Brunner et al. 2022).

### Conclusions

We revised the existing faunal list of Galápagos Rift vents and added 15 new records based on our observations and specimens collected, bringing the total to 92 species. Of these species, 14 are only known from Galápagos Rift. Though only based on qualitative observations, our results suggest some differences in fauna composition of vents at eastern vs western Galápagos Rift, warranting future research. Diversity data provide important grounds for constructing management strategies and spatial planning, especially with the growing interests for deep-sea mineral resources. As the Galápagos Rift is partially included in the Galápagos Marine Reserve, our updated species list will also be useful for conservation and marine spatial planning in this world heritage site. On one hand, we increase considerably the number of vent species (especially those living on high-temperature chimneys) living within the Reserve, including those lacking any formal protection on the extensive East Pacific Rise south of Mexican waters. On the other hand, we show that Galápagos Rift vents host several endemic species. This is further supplemented by cryptic lineages and genetic diversities not found outside Galápagos due to isolation from the EPR at least for some gastropods (Matabos and Jollivet 2019), likely also true for some newly recorded species herein. Altogether, our results highlight the Galápagos vents as a candidate for focused conservation efforts.

## Acknowledgements

We thank the captain and crew of R/V *Falkor (too)* during the research cruise Fkt231024 (‘Project Zombie: Bringing dead vents to life – Ultra fine-scale seafloor mapping’), as well as the ROV *SuBastian* team for their immense support. To clarify, (We Were Never Promised A) Rose Garden on this cruise. Viola Watts (University of Victoria), Janet Voight (The Field Museum), Hiromi K. Watanabe (JAMSTEC), and Anders Warén (Swedish Museum of Natural History) are thanked for helping with identifications and historical records. Ana-Belén Yánez Suárez (Marine Institute of Memorial University of Newfoundland) and Stuart Banks (Charles Darwin Research Station) are gratefully acknowledged for facilitating permissions to carry out research within the Galápagos Marine Reserve in Ecuador. Miwako Tsuda (JAMSTEC) helped us with molecular barcoding. The cruise FKt231024 was supported and authorised by the Galápagos National Park Directorate, the Instituto Oceanográfico y Antártico de la Armada de Ecuador (INOCAR) and facilitated by the Charles Darwin Foundation Deep-Ocean Research Program under permit number PC 51-23. This publication is contribution number 2602 of the Charles Darwin Foundation for the Galápagos Islands. We also thank our on-board Ecuadorian observers, Diego Bermeo (Galápagos National Park) and Richard Porfirio Narea Ortega (INOCAR). We recognize the extensive support from the Schmidt Ocean Institute for this expedition and associated logistics. The vent field name ‘Tortugas’ is a reference to the Galápagos giant tortoise (tortugas gigantes de las islas Galápagos, *Chelonoidis niger*) and was decided in consortia among the scientific party including discussions with the observers. Comments and edits from two anonymous reviewers helped improve an earlier version of this paper.

## Author Contributions

CC and VT conceived the project. CC and VT identified animal species, and updated the faunal list. CC undertook photography and made the figures. JWJ secured funding, planned and led the expedition, provided map of the research area. CC drafted the original manuscript which was critically revised by JWJ and VT. All authors agreed to the submission of the final version.

## Ethical approval

Study species were invertebrates and no experimental manipulation was undertaken on live animals in this study. All necessary permits for sampling and observation have been obtained by the authors where applicable. Permission for ROV dives inside the Galápagos Marine Reserve in Ecuador was granted through a partnership between Schmidt Ocean Institute and the Charles Darwin Foundation (MAATE-DPNG/DGA-2023-1449-O).

## Data availability

Data relevant to our conclusions are included in the main text. Specimens collected within the Galápagos Marine Reserve are vouchered in the Charles Darwin Research Station on Santa Cruz Island (https://www.darwinfoundation.org/en/about/cdrs). All ROV *SuBastian* dives during cruise FKt231024 are available online through the Schmidt Ocean Institute YouTube page: https://www.youtube.com/@SchmidtOcean/streams

## References

Anderson ME (1989) Review of the eelpout genus *Pachycara* Zugmayer, 1911 (Teleostei: Zoarcidae), with descriptions of six new species. Proceedings of the California Academy of Sciences 46: 221-242

Barnard JL, Ingram CL (1990) Lysianassoid Amphipoda (Crustacea) from deep-sea thermal vents. Smithsonian Contributions to Zoology 499: 1–80

Blake JA (1985) Polychaeta from the vicinity of deep-sea geothermal vents in the eastern Pacific. I: Euphrosinidae, Phyllodocidae, Hesionidae, Nereididae, Glyceridae, Dorvilleidae, Orbiniidae and Maldanidae. Bulletin of the Biological Society of Washington 6: 67–101

Borda E, Kudenov JD, Chevaldonné P, et al. (2013) Cryptic species of *Archinome* (Annelida: Amphinomida) from vents and seeps. Proceedings of the Royal Society B: Biological Sciences 280: 20131876 doi doi:10.1098/rspb.2013.1876

Boss KJ, Turner RD (1980) The giant white calm from the Galapagos Rift, *Calyptogena magnifica* species novum. Malacologia 20: 161–194

Brunner O, Chen C, Giguère T, et al. (2022) Species assemblage networks identify regional connectivity pathways among hydrothermal vents in the Northwest Pacific. Ecology and Evolution 12: e9612 doi 10.1002/ece3.9612

Burreson EM (1981) A new deep-sea leech, Bathybdella sawyeri, n. gen., n. sp., from thermal vent areas on the Galápagos Rift. Proceedings of the Biological Society of Washington 94: 483–491

Cavanaugh CM, Gardiner SL, Jones ML, Jannasch HW, Waterbury JB (1981) Prokaryotic Cells in the Hydrothermal Vent Tube Worm *Riftia pachyptila* Jones: Possible Chemoautotrophic Symbionts. Science 213: 340–342

Chen C, Ohara Y, Watanabe HK (2018) A very deep *Provanna* (Gastropoda: Abyssochrysoidea) discovered from the Shinkai Seep Field, Southern Mariana Forearc. Journal of the Marine Biological Association of the United Kingdom 98: 439–447 doi 10.1017/S0025315416001648

Chen C, Zhou Y, Watanabe HK, Zhang R, Wang C (2021) Neolepetopsid true limpets (Gastropoda: Patellogastropoda) from Indian Ocean hot vents shed light on relationships among genera. Zoological Journal of the Linnean Society 194: 276–296 doi 10.1093/zoolinnean/zlab081

Child CA (1987) *Ammothea verenae* and *Sericosura venticola*, two new hydrothermal vent associated pycnogonids from the Northeast Pacific. Proceedings of The Biological Society of Washington 100: 892–901

Child CA, Segonzac M (1996) *Sericosura heteroscela* And *S. cyrtoma*, New Species, And Other Pycnogonida From Atlantic And Pacific Hydrothermal Vents, With Notes On Habitat And Environment. Proceedings of the Biological Society of Washington 109: 664–676

Cohen DM, Rosenblatt RH, Moser HG (1990) Biology and description of a bythitid fish from deep-sea thermal vents in the tropical eastern Pacific. Deep Sea Research Part A Oceanographic Research Papers 37: 267–283 doi 10.1016/0198-0149(90)90127-H

Conroy-Dalton S, Huys R (1999) New genus of Aegisthidae (Copepoda: Harpacticoida) from hydrothermal vents on the Galapagos Rift. Journal of Crustacean Biology 19: 408–431

Corliss JB, Dymond J, Gordon LI, et al. (1979) Submarine Thermal Springs on the Galápagos Rift. Science 203: 1073–1083 doi doi:10.1126/science.203.4385.1073

de Saint Laurent M (1984) Crustacés Décapodes d’un site hydrothermal actif de la dorsale du Pacifique oriental (13° Nord), en provenance de la campagne française Biocyatherm. Comptes Rendus de l’Académie des Sciences 299: 355-360

de Saint Laurent M (1988) Les mégalopes et jeunes stades crabe de trois espèces du genre *Bythograea* Williams, 1980 (Crustacea Decapoda Brachyura). Oceanologica Acta Special Volume 8: 99-107

Desbruyères D, Laubier L (1982) *Paralvinella grasslei*, new genus, new species of Alvinellinae (Polychaeta: Ampharetidae) from the Galapagos Rift geothermal vents. Proceedings of the Biological Society of Washington 95: 484–494

Desbruyères D, Laubier L (1986) Les Alvinellidae, une famille nouvelle d’annélides polychètes inféodées aux sources hydrothermales sous-marines: systématique, biologie et écologie. Canadian Journal of Zoology 64: 2227–2245

Desbruyères D, Segonzac M, Bright M (2006) Handbook of Deep-sea Hydrothermal Vent Fauna (2nd Eds.). Denisia 18: 1-544

Fauchald K (1982) A Eunicid Polychaete From A White Smoker. Proceedings of the Biological Society of Washington 95: 781–787

Folmer O, Black M, Hoeh W, Lutz R, Vrijenhoek R (1994) DNA primers for amplification of mitochondrial cytochrome *c* oxidase subunit I from diverse metazoan invertebrates. Mol Mar Biol Biotechnol 3: 294–299

Giguère TN, Tunnicliffe V (2021) Beta diversity differs among hydrothermal vent systems: Implications for conservation. PLOS ONE 16: e0256637 doi 10.1371/journal.pone.0256637

Gollner S, Kaiser S, Menzel L, et al. (2017) Resilience of benthic deep-sea fauna to mining activities. Marine Environmental Research 129: 76–101 doi 10.1016/j.marenvres.2017.04.010

Guinot D, Hurtado A1 (2003) Two new species of hydrothermal vent crabs of the genus *Bythograea* from the southern East Pacific Rise and from the Galapagos Rift (Crustacea Decapoda Brachyura Bythograeidae). Comptes Rendus Biologies 326 423–439

Gustafson RG (1991) Distribution of molluscan morphospecies at deep-sea hydrothermal vents and cold-water seep areas. Abstracts of the 6th Deep-sea Biology Symposium, Copenhagen, 1991 1

Gustafson RG, Lutz RA (1994) Molluscan life history traits and deep-sea hydrothermal vents and cold methane/sulphide seeps. In: Young CM, Eckelbarger KJ (eds) Reproduction, Larval Biology, and Recruitment of the Deep-Sea Benthos. Columbia University Press, New York, pp 76–97

Haymon RM, White SM, Baker ET, et al. (2008) High-resolution surveys along the hot spot–affected Gálapagos Spreading Center: 3. Black smoker discoveries and the implications for geological controls on hydrothermal activity. Geochemistry, Geophysics, Geosystems 9 doi 10.1029/2008GC002114

Hessler RR (1984) *Dahlella caldariensis*, New Genus, New Species: A Leptostracan (Crustacea, Malacostraca) from Deep-Sea Hydrothermal Vents. Journal of Crustacean Biology 4: 655–664 doi 10.2307/1548079

Hessler RR, Smithey WM (1983) The Distribution and Community Structure of Megafauna at the Galapagos Rift Hydrothermal Vents. In: Rona PA, Boström K, Laubier L, Smith KL (eds) Hydrothermal Processes at Seafloor Spreading Centers. Springer US, Boston, MA, pp 735-770

Humes AG (1987) Copepods from deep-sea hydrothermal vents. Bulletin of Marine Science 41: 645–788

Humes AG (1988a) *Oncaea praeclara* n. sp. (Copepoda: Poecilostomatoida) from deep-sea hydrothermal vents in the eastern Pacific. Journal of Plankton Research 10: 475–485

Humes AG (1988b) *Hyalopontius boxshalli*, new species (Copepoda: Siphonostomatoida), from a deep-sea hydrothermal vent at the Galapagos Rift. Proceedings of the Biological Society of Washington 101: 825–831

Humes AG (1989) *Rhogobius pressulus* n. sp. (Copepoda, Siphonostomatoida) from a deep-sea hydrothermal vent at the Galapagos Rift. Pacific Science 43: 27–31

Humes AG, Dojiri M (1980) A siphonostome copepod associated with a vestimentiferan from the Galapagos Rift and the East Pacific Rise. Proceedings of the Biological Society of Washington 93: 697–707

Humes AG, Segonzac M (1998) Copepoda from deep-sea hydrothermal sites and cold seeps: Description of a new species of *Aphotopontius* from the East Pacific Rise and general distribution. Cahiers de Biologie Marine 39: 51–62

Hurtado LA, Lutz RA, Vrijenhoek RC (2004) Distinct patterns of genetic differentiation among annelids of eastern Pacific hydrothermal vents. Molecular Ecology 13: 2603–2615 doi 10.1111/j.1365-294X.2004.02287.x

Jones ML (1981) *Riftia pachyptila*, new genus, new species, the vestimentiferan worm from the Galápagos Rift geothermal vents (Pogonophora). Proceedings of the Biological Society of Washington 93: 1295–1313

Jones WJ, Macpherson E (2007) Molecular Phylogeny of the East Pacific Squat Lobsters of the Genus *Munidopsis* (Decapoda: Galatheidae) with the Descriptions of Seven New Species. Journal of Crustacean Biology 27: 477–501 doi 10.1651/s-2791.1

Kenk VC, Wilson BR (1985) A new mussel (Bivalvia, Mytilidae) from hydrothermal vents, in the Galapagos Rift zone. Malacologia 26: 253–271

Komai T, Tsuchida S, Michel S (2012) Records of species of the hippolytid genus *Lebbeus* White, 1847 (Crustacea: Decapoda: Caridea) from hydrothermal vents in the Pacific Ocean, with descriptions of three new species. Zootaxa 3241: 35-63 doi 10.11646/zootaxa.3241.1.2

Krantz GW (1982) A new species of *Copidognathus* Trouessart (Acari: Actinedida: Halacaridae) from the Galapagos Rift. Canadian Journal of Zoology 60: 1728–1731 doi 10.1139/z82-225

Maciolek NJ (1981) Spionidae (Annelida: Polychaeta) from the Galapagos Rift geothermal vents. Proceedings of the Biological Society of Washington 94: 826–837

Matabos M, Jollivet D (2019) Revisiting the *Lepetodrilus elevatus* species complex (Vetigastropoda: Lepetodrilidae), using samples from the Galápagos and Guaymas hydrothermal vent systems. Journal of Molluscan Studies 85: 154–165 doi 10.1093/mollus/eyy061

McLean JH (1981) The Galapagos Rift limpet *Neomphalus*: relevance to understanding the evolution of a major Paleozoic-Mesozoic radiation. Malacologia 21: 291–336

McLean JH (1988) New archaeogastropod limpets from hydrothermal vents; superfamily lepetodrilacea I. Systematic descriptions. Philosophical Transactions of the Royal Society of London B, Biological Sciences 319: 1–32 doi doi:10.1098/rstb.1988.0031

McLean JH (1989a) New archaeogastropod limpets from hydrothermal vents: new family Peltospiridae, new superfamily Peltospiracea. Zoologica Scripta 18: 49–66 doi 10.1111/j.1463-6409.1989.tb00123.x

McLean JH (1989b) New slit-limpets (Scissurellacea and Fissurellacea) from hydrothermal vents. Part 1. Systematic descriptions and comparisons based on shell and radular characters. Contributions in science 407: 1-29 doi 10.5962/p.208131

McLean JH (1990) Neolepetopsidae, a new docoglossate limpet family from hydrothermal vents and its relevance to patellogastropod evolution. Journal of Zoology 222: 485–528 doi 10.1111/j.1469-7998.1990.tb04047.x

Moalic Y, Desbruyères D, Duarte CM, et al. (2012) Biogeography Revisited with Network Theory: Retracing the History of Hydrothermal Vent Communities. Systematic Biology 61: 127–127 doi 10.1093/sysbio/syr088

Pettibone MH (1984a) A new scale-worm commensal with deep-sea mussels on the Galapagos hydrothermal vent (Polychaeta: Polynoidae). Proceedings of the Biological Society of Washington 97: 226–239

Pettibone MH (1984b) Two new species of *Lepidonotopodium* (Polychaeta: Polynoidae: Lepidonotopodinae) from hydrothermal vents off the Galapagos and East Pacific Rise at 21 N. Proceedings of the Biological Society of Washington 97: 849–863

Pettibone MH (1985a) Additional branchiate scale-worms (Polychaeta: Polynoidae) from Galapagos hydrothermal vent and rift-area off western Mexico at 21 N. Proceedings of the Biological Society of Washington 98: 447–469

Pettibone MH (1985b) New genera and species of deep-sea Macellicephalinae and Harmothoinae (Polychaeta: Polynoidae) from the hydrothermal rift areas off the Galapagos and western Mexico at 21°N and from Santa Catalina Channel. Proceedings of the Biological Society of Washington 98: 740–757

Pettibone MH (1990) New species and new records of scaled polychaetes (Polychaeta: Polynoidae) from the Axial Seamount Caldera of the Juan de Fuca ridge in the northeast Pacific Ocean off northern California. Proceedings of the Biological Society of Washington 103: 825–838

Pleijel F, Rouse GW, Nygren A (2012) A revision of *Nereimyra* (Psamathini, Hesionidae, Aciculata, Annelida). Zoological Journal of the Linnean Society 164: 36–51 doi 10.1111/j.1096-3642.2011.00756.x

Pugh PR, Marshall NB (1983) Benthic siphonophores: a review of the family Rhodaliidae (Siphonophora, Physonectae). Philosophical Transactions of the Royal Society of London B, Biological Sciences 301: 165–300 doi doi:10.1098/rstb.1983.0025

Raineault N, Ballard R, Mayer L, et al. (2016) Exploration of hydrothermal vents Along the Galápagos Spreading Center. Oceanography 29: 35–37

Rosenblatt RH, Cohen DM (1986) Fishes living in deepsea thermal vents in the tropical eastern Pacific, with descriptions of a new genus and two new species of eelpouts (Zoarcidae). Transactions of the San Diego Society of Natural History 21: 71–79

Salinas-de-León P, Phillips B, Ebert D, et al. (2018) Deep-sea hydrothermal vents as natural egg-case incubators at the Galapagos Rift. Scientific Reports 8: 1788 doi 10.1038/s41598-018-20046-4

Schein-Fatton E (1985) Découverte sur la ride du Pacifique oriental à 13°N d’un Pectinidae (Bivalvia, Pteriomorphia) d’affinités paléozoïques. Comptes Rendus de l’Academie des Sciences de Paris, Ser 3 301: 491–496

Scheltema A (2000) Two new hydrothermal vent species, *Helicoradomenia bisquama* and *Helicoradomenia acredema*, from the eastern Pacific Ocean (Mollusca, Aplacophora). Argonauta 14: 15–25

Schmidt Ocean Institute (2023) Sendero del Cangrejo | SOI Divestream 573 YouTube, Schmidt Ocean Channel, https://youtu.be/5rlniuALVnM?t=23684

Sclater JG, Klitgord KD (1973) A detailed heat flow, topographic, and magnetic survey across the Galapagos Spreading Center at 86°W. Journal of Geophysical Research (1896–1977) 78: 6951-6975 doi 10.1029/JB078i029p06951

Segonzac M, Vervoort W (1995) First record of the genus *Candelabrum* (Cnidaria, Hydrozoa, Athecata) from the Mid-Atlantic Ridge: a description of a new species and a review of the genus. Bulletin du Muséum national d’histoire naturelle 17: 31–63 doi 10.5962/p.290312

Shank T, Baker E, Embley R, et al. (2012) Exploration of the deepwater Galápagos region. Oceanography 25: 50–51

Shank T, Fornari D, Yoerger D, et al. (2003) Deep submergence synergy: Alvin and ABE explore the Galapagos Rift at 86°W. Eos, Transactions American Geophysical Union 84: 425–433 doi 10.1029/2003EO410001

Ten Hove HA, Zibrowius H (1986) *Laminatubus alvini* gen. et sp. n. and *Protis hydrothermica* sp. n. (Polychaeta, Serpulidae) from the bathyal hydrothermal vent communities in the eastern Pacific. Zoologica Scripta 15: 21–31

Tunnicliffe V (1991) The biology of hydrothermal vents: Ecology and evolution. Marine Biology and Oceanography: An Annual Review 29: 319–407

Tunnicliffe V (1992) The Nature and Origin of the Modern Hydrothermal Vent Fauna. PALAIOS 7: 338–350 doi 10.2307/3514820

Tunnicliffe V, Chen C, Giguère T, et al. (2023) Hydrothermal vent fauna of the western Pacific Ocean: Distribution patterns and biogeographic networks. Diversity and Distributions doi 10.1111/ddi.13794

Tunnicliffe V, Juniper SK, Burgh MED (1985) The hydrothermal vent community on axial seamount, Juan de Fuca ridge. Bulletin of The Biological Society of Washington: 453–464

Tyler PA, Pendlebury S, Mills SW, et al. (2008) Reproduction of Gastropods from Vents on the East Pacific Rise and the Mid-Atlantic Ridge. Journal of Shellfish Research 27: 107–118, 112

Wang J, Lin R, Bamber R, Dingyong H (2013) Two new species of *Sericosura* Fry & Hedgpeth, 1969 (Arthropoda: Pycnogonida: Ammotheidae) from a hydrothermal vent on the East Pacific Rise. Zootaxa 3669: 165–171 doi 10.11646/zootaxa.3669.2.8

Warén A, Bouchet P (1986) Four new species of *Provanna* Dall (Prosobranchia, Cerithiacea?) from East Pacific hydrothermal sites. Zoologica Scripta 15: 157–164 doi 10.1111/j.1463-6409.1986.tb00218.x

Warén A, Bouchet P (1989) New gastropods from East Pacific hydrothermal vents. Zoologica Scripta 18: 67–102 doi 10.1111/j.1463-6409.1989.tb00124.x

Warén A, Bouchet P (1993) New records, species, genera, and a new family of gastropods from hydrothermal vents and hydrocarbon seeps. Zoologica Scripta 22: 1–90 doi 10.1111/j.1463-6409.1993.tb00342.x

Warén A, Bouchet P (2001) Gastropoda and Monoplacophora from hydrothermal vents and seeps; New taxa and records. The Veliger 44: 116–231

Weiss RF, Lonsdale P, Lupton JE, Bainbridge AE, Craig H (1977) Hydrothermal plumes in the Galapagos Rift. Nature 267: 600–603 doi 10.1038/267600a0

Williams AB (1980) A new crab family from the vicinity of submarine thermal vents on the Galapagos Rift (Crustacea: Decapoda: Brachyura). Proceedings of the Biological Society of Washington 93: 443–472

Williams AB, Chace FAJ (1982) A new caridean shrimp of the family Bresiliidae from thermal vents of the Galapagos Rift. Journal of Crustacean Biology 2: 136–147

Woodwick K, Sensenbaugh T (1985) *Saxipendium coronatum*, new genus, new species (Hemichordata: Enteropneusta): the unusual spaghetti worms of the Galápagos Rift hydrothermal vents. Proceedings of the Biological Society of Washington 98: 351–365

WoRMS Editorial Board (2023) World Register of Marine Species. Available from https://www.marinespecies.org/ at VLIZ Accessed 2023-11-14: DOI: 10.14284/14170

Zottoli R (1983) *Amphisamytha galapagensis* a new species of ampharetid polychaete from the vicinity of abyssal hydrothermal vents in the Galapagos Rift, and the role of this species in rift ecosystems. Proceedings of the Biological Society of Washington 96: 379–391

